# Mixed-mobility supported lipid bilayers reveal the modulatory role of immobilized ICAM1 on T cell activation, effector functions and immune synapse organization

**DOI:** 10.1101/2025.09.12.675755

**Authors:** Alexander Leithner, Audun Kvalvaag, Tanmay Mitra, Salvatore Valvo, Hannah Dada, Ewoud Compeer, Edward Jenkins, Christoffer Lagerholm, Omer Dushek, Michael L. Dustin

**Author notes:** contributed equally (Audun Kvalvaag, Tanmay Mitra, Salvatore Valvo).

## Abstract

The immunological synapse (IS) integrates antigen recognition and adhesion to control T cell activation and effector functions. Reductionist systems have been instrumental in dissecting IS organization, but conventional systems constrain all ligands to be either mobile or immobile, unlike antigen-presenting cells where intercellular adhesion molecule 1 (ICAM1) is cytoskeletally anchored while T cell receptor (TCR) ligands remain mobile. Here, we establish mixed-mobility supported lipid bilayers (SLBs) that simultaneously present mobile TCR agonists and immobile ICAM1. Selective immobilization of ICAM1 disrupts centripetal F-actin flow, prevents centralization of TCR microclusters and shifts signaling to peripheral microclusters. This attenuates TCR downregulation through ectocytosis while maintaining recycling, and enhances integrin mechanotransduction, reflected in increased phosphorylation of Focal Adhesion Kinase, Paxillin and the stretch-sensitive adaptor CasL. Functionally, immobilized ICAM1 augments T cell activation, degranulation, Perforin release and cytotoxicity. Importantly, these findings were recapitulated in a cell–cell system engineered to express either full-length, cytoskeleton-anchored ICAM1 or a truncated form lacking cytoskeletal association, with full-length ICAM1 consistently promoting stronger effector responses. These findings identify ligand mobility as a key biophysical parameter that shapes IS organization and T cell effector responses and establish mixed-mobility SLBs as a powerful tool for probing receptor mechanics in immunity.

## Introduction

The immunological synapse (IS) is a highly organized cell-cell interface that governs critical T cell functions, including antigen recognition, signal integration and effector responses (Friedl, den Boer, and Gunzer 2005; Norcross 1984). Mechanistic insights into IS formation have largely emerged from a number of reductionist *in vitro* systems. Initial seminal studies with cell lines serving as model antigen presenting cells (APCs) first demonstrated formation of the central supramolecular activation cluster (cSMAC), enriched for T cell receptor (TCR) and protein kinase C-Θ (PKC-Θ), surrounded by the peripheral SMAC (pSMAC) enriched for lymphocyte function-associated antigen 1 (LFA-1), talin and F-actin (Monks et al. 1998; Krummel et al. 2000). Simple glass-supported lipid bilayers (SLBs), presenting laterally mobile peptide Major Histocompatibility Complex (pMHC) or TCR agonist and Intercellular adhesion molecule 1 (ICAM1) generated similar IS dynamics and organisation, while enabling quantitative, high-resolution microscopy (Grakoui et al. 1999; Yokosuka et al. 2005; Somersalo et al. 2004; Yokosuka et al. 2008). In parallel, glass substrates coated with immobile stimulatory antibodies (Bunnell et al. 2002) provided insights into microclusters, the basic TCR signalling unit, which were also glimpsed with model APCs (Krummel et al. 2000) and extensively studied in SLBs (Varma et al. 2006).

However, while enabling valuable insights into the monofocal IS, each of these model systems carries certain inherent limitations. The employed model APCs lack active cytoskeletal processes that are present in professional APCs (Al-Alwan et al. 2001; Wülfing and Davis 1998), which form multifocal instead of monofocal synapses (Tseng et al. 2008; Brossard et al. 2005; Leithner et al. 2021; Fisher et al. 2008). Crucially, a limitation of the SLB system is that ligands like ICAM1 are presented in a laterally mobile form, whereas natural ICAM1 can be anchored to the cytoskeleton (Carpén et al. 1992; Barreiro et al. 2002). In this form, ICAM1 can resist the mechanical forces transmitted through LFA-1, which is connected via talin to the actin cytoskeleton of the T cell (Nordenfelt et al. 2017). Conversely, solid phase presentation has the limitation that all ligands are immobilised, which is not the normal state of most membrane proteins, including MHC, which are laterally mobile (Bierer et al. 1987; Schlessinger et al. 1976). Similarly, variations of the SLB system that contain nanofabricated chrome barriers or frozen lipids can alter ligand mobility but again impact all ligands in the SLB in the same manner (Mossman et al. 2005; Hsu et al. 2012). Crucially, in dendritic cells (DCs), anchorage of ICAM1 to the F-actin cytoskeleton via its cytoplasmic tail, alongside the simultaneous presentation of laterally mobile MHCII, is associated with optimal T cell responses (Comrie et al. 2015).

This observation supports the hypothesis that selective immobilization of ICAM1 enhances T cell activation, highlighting the need for new methodologies that enable the simultaenous presentation of both mobile and immobile ligands. To directly test how ICAM1 anchorage affects T cell IS formation, activation and degranulation, we developed a *mixed-mobility* SLB platform enabling independent control over the lateral mobility of anti-TCR and ICAM1. Using this system, we demonstrate that selective ICAM1 immobilization profoundly influences multiple aspects of IS formation, including T cell spreading dynamics, F-actin cytoskeleton organization, TCR microcluster transport, TCR recycling and integrin activation. Importantly, we show that ICAM1 immobilization enhances T cell degranulation and Perforin-1 release, findings further supported by increased cytotoxicity in cell–cell killing assays.

Together, these results highlight the importance of mobility regulation as a key biophysical parameter that shapes T cell responses and emphasize its relevance for both mechanistic studies and the design of future immunotherapies.

## Results

To establish a reductionist system that enables the simultaneous presentation of laterally mobile and immobile ligands to T cell-expressed receptors, we built on previous findings showing that, in SLBs formed from proteoliposomes containing type I transmembrane proteins with large extracellular domains (ECDs) and relatively short cytoplasmic tails, the ECD is exposed to the medium while the cytoplasmic domain becomes trapped against the glass surface, rendering these proteins laterally immobile (Dustin et al. 1996; McConnell et al. 1986; Chan et al. 1991). We hypothesized that reconstitution of full-length, transmembrane ICAM1 (ICAM1-FL) into nickel-nitrilotriacetic acid (Ni-NTA) - containing liposomes would allow the formation of *mixed-mobility* SLBs on glass surfaces, presenting immobile, glass-anchored ICAM1 alongside mobile, histidine (His)-tagged ligands, in this case the anti-TCR Fab (**Fig.1A**). In order to accomplish this, we first had to purify ICAM1-FL.

ICAM1-FL was solubilized from human spleen tissue in 1% Tritox X-100 detergent, captured on anti-ICAM1 functionalized Agarose beads that were extensively washed, followed by exchange of the detergent to dialyzable 1% n-octyl-β-D-glucopyranoside. ICAM1 was then eluted at pH 3, neutralized and subsequently reconstituted into 1,2-dioleoyl-sn-glycero-3-phosphocholine (DOPC) proteoliposomes by dialysis as previously described (Dustin and Springer 1988) with the addition that we also included 2 mol% of the Ni^2+^ salt of 1,2-dioleoyl-sn-glycero-3-[(N-(5-amino-1-carboxypentyl)iminodiaceticacid)succinyl (DOGS-NTA). Liposomes of the same composition, but lacking ICAM1-FL, served as controls. The ICAM1-FL density in each batch of proteoliposome preparation was determined by forming SLBs on glass beads, followed by staining with a fluorescently labeled anti-ICAM1 antibody of known fluorophore-to-protein (F/P) ratio and quantitative flow cytometry. The resulting densities ranged from 50 to 200 molecules/µm^2^. SLBs from proteoliposomes and control liposomes were then formed on glass and further functionalized with His-tagged, Alexa Fluor 647–labeled anti-TCR Fab fragments at a density of 30 molecules/µm^2^. Additionally, SLBs formed from control liposomes were loaded with His-tagged ICAM1-ECD to match the predetermined surface densities of ICAM1-FL in the proteoliposome-derived SLBs. ICAM1-FL and ICAM1-ECD on the SLBs were then labelled with an Alexa Fluor 488–conjugated anti-ICAM1 antibody, and fluorescence recovery after photobleaching (FRAP) experiments were performed to assess their lateral mobility.

ICAM1-FL and ICAM1-ECD on the SLB were labelled with an Alexa Fluor 488–conjugated anti-ICAM1 antibody, and fluorescence recovery after photobleaching (FRAP) experiments were performed to assess their lateral mobility.

In both conditions, His-tagged anti-TCR Fab fragments displayed rapid recovery (τ ∼ 4 s for a spot radius of 2µm), with a mobile fraction of almost 100% (**Fig. 1B - D**). Similarly, ICAM1-ECD exhibited rapid (τ ∼ 6 s) and nearly complete recovery after correction for photobleaching, indicating free lateral mobility, as expected. In contrast, ICAM1-FL displayed slow recovery (τ >> 1 m) with >95% immobile over 1 min. (**Fig. 1B-D**). To further investigate potential effects of immobile ICAM1 on the mobility of anti-TCR Fab, we performed point-scan fluorescence correlation spectroscopy (FCS) measurements. The mean diffusion coefficients of anti-TCR Fab fragments, in the presence of either ICAM1-FL or ICAM1-ECD, were indistinguishable and averaged around 1 µm^2^/sec. (**Fig. 1E**), consistent with previous measurements for fully mobile proteins in SLBs. Thus, the presence of immobile ICAM1-FL does not impact the mobility of other proteins that interact with the upper leaflet of the SLB.

**Figure 1:**
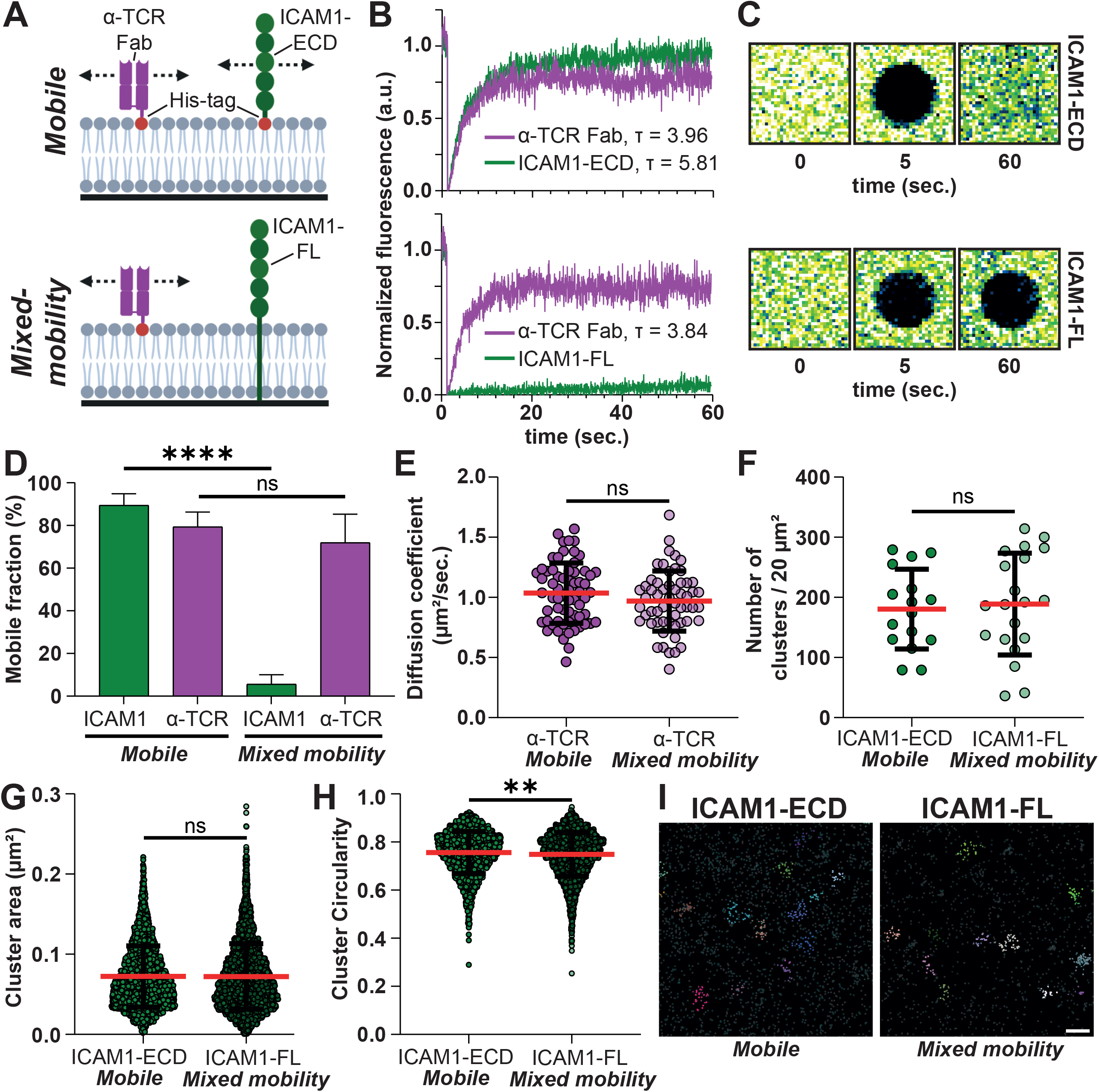
(**A**) Schematic overview over the mobile (top) and *mixed-mobility* (bottom) SLB system. (**B**) FRAP measurements of fluorescently labelled α-TCR Fab and ICAM1-ECD (top) or α-TCR Fab and ICAM1-FL (bottom). (**C**) Representative example images of fluorescence recovery of ICAM1-ECD (top) and ICAM1-FL (bottom). (**D**) Quantification of the mobile fraction of ICAM1 and α-TCR Fab in the mobile and *mixed-mobility* SLB. Unpaired t-tests, **** <0.0001. (**E**) Quantification of diffusion coefficients of α-TCR Fab in the context of mobile and *mixed-mobility* SLBs. Unpaired t-test. (**F**) Quantification of the number of ICAM1-ECD and ICAM1-FL clusters per 20µm^2^. Unpaired t-test. (**G**) Quantification of the area of ICAM1-ECD and ICAM1-FL clusters in µm^2^. Unpaired t-test. (**H**) Quantification of the circularity of ICAM1-ECD and ICAM1-FL clusters. Unpaired t-test, **<0.01. (**I**) Representative images of ICAM1 clusters identified by dSTORM.

Additionally, we performed direct Stochastic Optical Reconstruction Microscopy (dSTORM) to characterize the nanoscale organisation of ICAM1-ECD and ICAM1-FL in SLBs. Data acquisition was followed by hierarchical density-based clustering (Campello, Moulavi, and Sander 2013) and cluster analysis. Nanoscale clusters were identified for both conditions, ICAM1-ECD and ICAM1-FL, whose formation was potentially mediated by oligomerization of their extracellular domains (Yang et al. 2004). No significant differences could be observed in protein cluster density (**Fig. 1F**) or cluster area (**Fig. 1G**). We only detected a very small but statistically significant difference in mean protein cluster circularity, from ∼0.76 for ICAM1-ECD to ∼0.75 for ICAM1-FL (**Fig. 1H**), indicating a slightly different shape between ICAM1-ECD and ICAM1-FL protein clusters. These findings suggest that T cells would initially encounter a comparable molecular distribution of ICAM1 on either surface.

Taken together, our data demonstrate that reconstituting a mixture of full-length transmembrane proteins and His-tagged ECDs on Ni^2+^–NTA–functionalized SLBs enables the formation of *mixed-mobility* SLBs. From here on, we refer to the SLB containing ICAM1-FL and His-tagged anti-TCR Fab as the *mixed-mobility* SLB, and to the control surface containing His-tagged ICAM1-ECD and anti-TCR Fab as the *mobile* SLB.

Next, we sought to study the functional effects of selective ICAM1 immobilisation on T cell activation by mobile TCR ligands. To this end, we introduced CD8^+^ T cells to the *mobile* and *mixed-mobility* SLBs and allowed them to interact for 3h. The cells were then removed and incubated for another 18h, after which surface expression of the T cell activation marker CD69 was assessed, consistent with the notion that T cells require a short period of stimulation to execute functional programs (van Stipdonk, Lemmens, and Schoenberger 2001). While T cells stimulated on *mobile* SLBs only exhibited low levels of CD69 upregulation, stimulation on *mixed-mobility* SLBs resulted in significantly higher CD69 expression, suggesting that immobile ICAM1 is a more active co-stimulator than laterally mobile ICAM1 (**Fig. 2A**).

**Figure 2:**
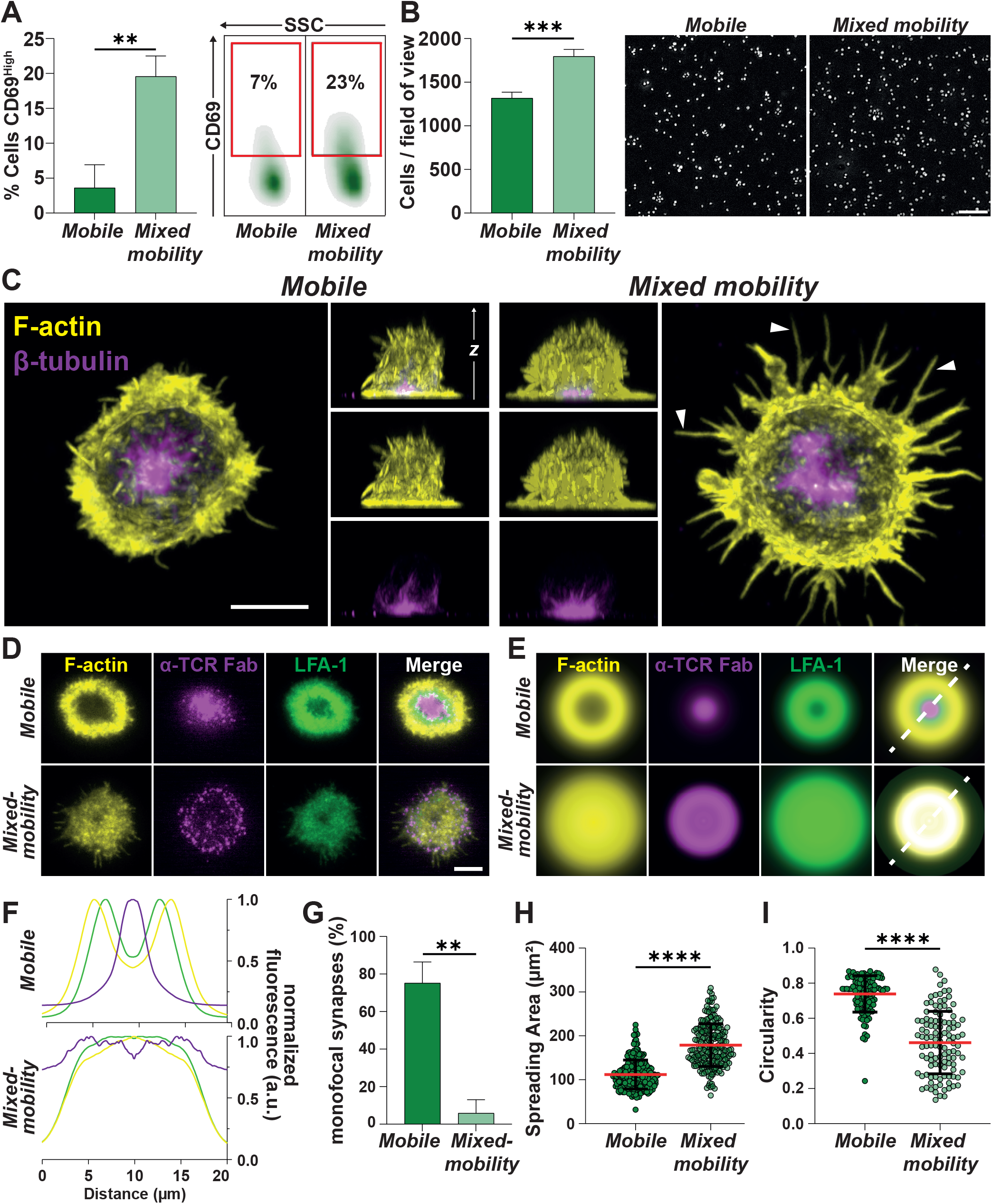
(**A**) Quantification (left) and representative example (right) of flow cytometry stainings for CD69^High^ T cells after 3h on mobile and *mixed-mobility* SLBs and subsequent incubation for 18h. Unpaired t-test, **<0.01. 3 Biological replicates. (**B**) Quantification (left) and representative example images (right) of cells per field of view on mobile and *mixed-mobility* SLBs after 15min. of incubation. Unpaired t-test, *** = 0.0001. 4 Biological replicates. Scale bar = 20µm. (**C**) Representative maximum intensity projections and orthogonal views of Airyscan confocal microscopy z-stacks of T cells seeded onto mobile and *mixed-mobility* SLBs, fixed and permeabilized after 15min. of interaction and stained with Phalloidin (F-actin, yellow) and a directly labelled antibody for β-tubulin (magenta). Scale bar = 5µm. White arrows demarcate filopodial projections on *mixed-mobility* SLBs. (**D**) Representative TIRF images of synapses formed by T cells on mobile and *mixed-mobility* SLBs, containing fluorescently labelled α-TCR Fab (magenta) after 15min. of incubation. Cells were fixed, stained with a directly labelled antibody for LFA-1 (green) and permeabilized and stained with Phalloidin (F-actin, yellow). Scale bar = 5µm. (**E**) Average intensity projections of radial averages of ≥56 individual synapses formed on mobile or *mixed-mobility* SLBs. (**F**) Normalized intensity profiles along white dashed lines in (E) for all labelled proteins. (**G**) Quantification of monofocal synapses on mobile and *mixed-mobility* SLBs. Mann-Whitney test, ** = 0.0022. 6 Biological replicates. (**H**) Quantification of the spreading area in µm^2^ and its circularity (**I**) of T cells spreading on mobile or *mixed-mobility* SLBs. Mann-Whitney test, **** < 0.0001. 3 Biological replicates.

In order to begin to understand the underlying cell biological principles of this effect, we first quantified the number of attached cells after 15 minutes of incubation and subsequent fixation. This revealed a moderate but significant effect on overall cell adhesion with more T cells attaching to *mixed-mobility* SLBs (**Fig. 2B**). Airy-Scan^®^ confocal microscopy of fixed, β-tubulin-stained T cells showed centrosome polarization towards the interaction plane under both mobile and *mixed-mobility* SLBs, suggesting that centrosome docking, a key step in IS formation(Tsun et al. 2011), remained unaffected (**Fig. 2C**). However, staining with fluorescently labelled phalloidin for F-actin revealed that T cells on *mixed-mobility* SLBs exhibited a different spreading morphology characterized by numerous filopodial projections not observed to the same extent on *mobile* SLBs (**Fig. 2C**).

Next, we used total internal reflection (TIRF) microscopy to assess the organization of the T cell-SLB contact area in more detail. As expected, T cells on *mobile* SLBs formed canonical synapses with high frequencies, characterized by concentric rings of intense F-actin and LFA-1 and a central accumulation of anti-TCR Fab, the canonical monofocal IS. In stark contrast, T cells on *mixed-mobility* SLBs primarily formed multifocal ISs in which F-actin and LFA-1 were uniformly distributed throughout the synaptic interface (**Fig. 2D-G**). Strikingly, anti-TCR Fab was not centralized but largely remained in the periphery as multiple discrete, bright microclusters (**Fig. 2D**). Furthermore, T cells on *mixed-mobility* SLBs exhibited a significantly increased spreading area, with the aforementioned projections leading to a decrease in circularity (**Fig. 2H & I**).

Centripetal flow of F-actin through classic dendritic actin nucleation, cofilin-dependent depolymerisation in the distal compartment and myosin-based contractility of anti-parallel F-actin bundles drives centralization of the TCR on mobile SLBs (Babich et al. 2012; Kaizuka et al. 2007; Murugesan et al. 2016). Additionally, integrin engagement by immobile ligands linked to solid substrates has been shown to slow down F-actin flow in IS formed on anti-TCR coated glass (Jankowska et al. 2018; Nguyen, Sylvain, and Bunnell 2008). Therefore, we aimed to assess how immobilization of ICAM1 in *mixed-mobility* SLBs influences F-actin flow and TCR movement. To this end, we introduced Lifeact-mCitrine mRNA into T cells by electroporation, seeded the cells onto SLBs and monitored the first two minutes of IS formation via time-lapse TIRF microscopy. In both SLB types, *mobile* and *mixed-mobility*, IS formation began with F-actin foci developing at initial contacts that rapidly expanded through an F-actin rich lamellipodium (**Fig. 3A & B, Supplementary Movie 1**). In the case of *mobile* SLBs, initial spreading was accompanied by the appearance of a central region with lower F-actin, a contraction of the overall contact, establishment of the F-actin transport network and the accumulation of anti-TCR Fab clusters at the centre of the IS (**Fig. 3A**). On the *mixed-mobility* SLB, the contrast between the initial bright F-actin ring and the interior of the IS was even more pronounced. However, unlike the situation on *mobile* SLBs, the F-actin signal reappeared in the central region by ∼ 90 s, accompanied by limited contraction, and no uniform centripetal F-actin flow (**Fig. 3B, Supplementary Movie 1 & 2**). Instead, the F-actin network exhibited fluctuations that sometimes appeared to propagate as waves (**Fig. 3C, Supplementary Movie 2**). Importantly, while anti-TCR Fab clusters on mobile SLB exhibited mean flow speeds of ∼ 0.05 µm/s and demonstrated considerable directionality consistent with centripetal flow, they remained relatively static after their initial formation on mixed-mobility SLB, showing significantly reduced speed (**Fig. 3E**) and directionality compared to *mobile* SLBs (**Fig. 3F & G**).

**Figure 3:**
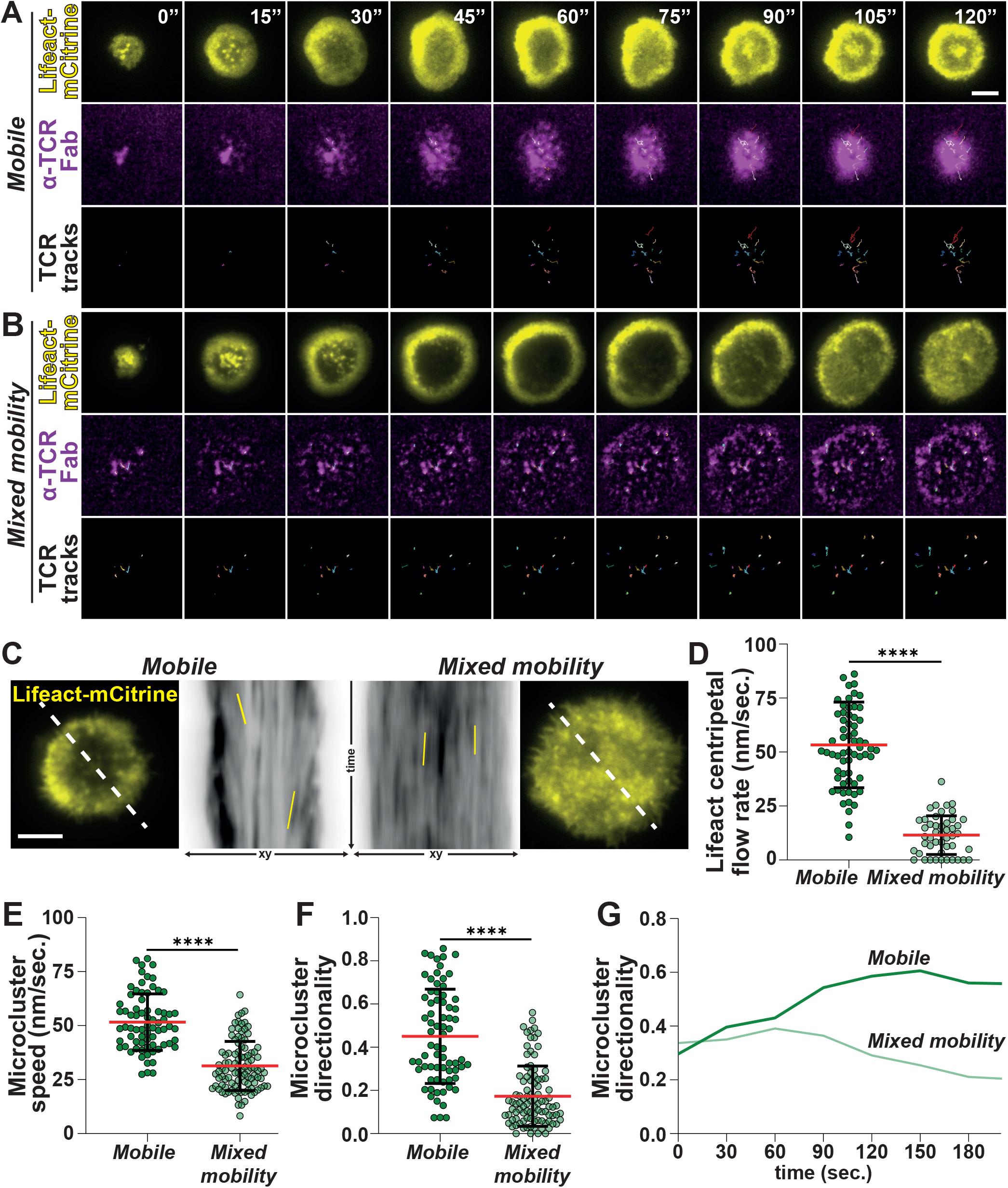
(**A**) Time-lapse TIRF microscopy of representative examples of Lifeact-mCitrine (yellow, top) transfected T cells interacting with mobile or (**B**) *mixed-mobility* SLBs containing fluorescently labelled α-TCR Fab (magenta, middle). Individual α-TCR Fab tracks are shown in colour on black background (bottom). Scale bar = 5µm. (**C**) Representative example images of time-lapse microscopy of Lifeact-mCitrine (yellow) transfected T cells interacting with mobile or *mixed-mobility* SLBs (outsides). White dashed lines were used for kymograph analysis (centre). Solid yellow lines demarcate examples of F-actin features that were used to determine the centripetal flow rate in nm/sec. shown in (**D**) Unpaired t-test, **** < 0.0001. Circles represent individual measurements from ≥10 individual cells per condition, 2 Biological replicates. (**E**) Quantification of microcluster speed in nm/sec. and directionality (**F**). Mann-Whitney test, **** < 0.0001. Circles represent individual tracks from 4 different cells per condition, 2 Biological replicates. (**G**) Average directionality of microclusters from (F) over time.

Taken together, these data suggest that engagement of immobile ICAM1 by T cell LFA-1 interferes with the formation of a centripetal F-actin flow, which is required for TCR centralization.

These findings led us to investigate how the multifocal IS might affect TCR downregulation and recycling. These processes have been originally linked to the formation of the cSMAC (Varma et al. 2006), where a biphasic process occurs: first, TCR-loaded vesicles are released from the plasma membrane, followed by TCR internalization (Kvalvaag et al. 2023). Upon TCR triggering, the receptor is ubiquitinated and recognized by Hepatocyte Growth Factor-Regulated Tyrosine Kinase Substrate (HRS), a component of the Endosomal Sorting Complex Required for Transport (ESCRT). HRS recruits clathrin and mediates the budding of TCR-loaded ectosomes from the plasma membrane. In the second phase, HRS is replaced by the endocytic adaptor protein Epsin-1 (EPN1), which promotes clathrin-dependent endocytosis. While the ectocytic pathway leads to TCR downregulation, EPN1-dependent internalization can either route the TCR for degradation or enable its recycling back to the plasma membrane. This model, originally developed based on data from *mobile* SLBs, has also been validated in cell-cell synapses (Kvalvaag et al. 2023).

Consistent with this model, T cells interacting with *mobile* SLBs exhibited increasing recruitment of anti-TCR Fab and high colocalization with HRS at early time points, which gradually declined over the course of the experiment, along with overall HRS levels (**Fig. 4A, C, D & F**). In parallel, initially low levels of EPN1 progressively increased, as did its colocalization with anti-TCR Fab, mostly in a central cluster (**Fig. 4E & G**). In contrast, under *mixed-mobility* conditions, the levels of anti-TCR Fab, HRS, and EPN1 remained relatively stable over time, with significantly lower levels of anti-TCR Fab and HRS compared to *mobile* SLBs (**Fig. 4B-E**). Colocalization coefficients for HRS/anti-TCR Fab remained comparatively low and unchanged throughout the time course (**Fig. 4F**). In contrast, colocalization between EPN1/anti-TCR Fab, although lower than on *mobile* SLBs, increased significantly over time (**Fig. 4G**).

**Figure 4:**
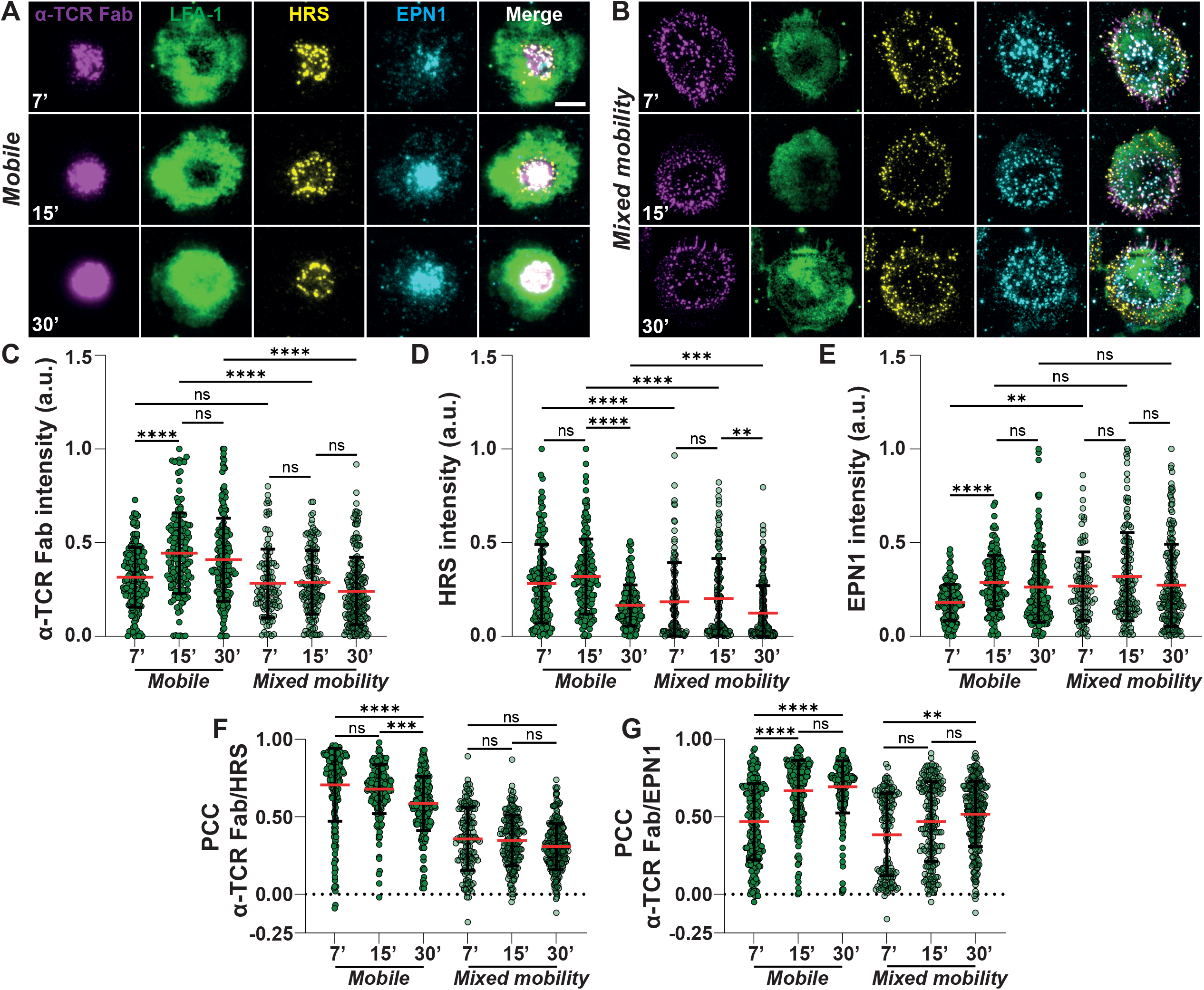
(**A**) Representative TIRF images of synapses formed by T cells on mobile and *mixed-mobility* SLBs (**B**), containing fluorescently labelled α-TCR Fab (magenta) after 7 (top), 15 (middle) and 30min (bottom). of incubation. Cells were fixed, stained with a directly labelled antibody for LFA-1 (green), permeabilized and stained with directly labelled antibodies for HRS (yellow) and EPN1 (cyan). Scale bar = 5µm. (**C**) Quantification of normalized α-TCR Fab, HRS (**D**) and EPN1 (**E**) intensities after indicated times of incubation on mobile and *mixed-mobility* SLBs. Kruskal-Wallis test, ** < 0.01, *** < 0.001, **** < 0.0001. Circles represent measurements from individual cells. 3 Biological replicates. (**F**) Pearson correlation coefficients of α-TCR Fab and HRS or α-TCR Fab and EPN1 (**G**), respectively. Kruskal-Wallis test, ** < 0.01, *** < 0.001, **** < 0.0001. 3 Biological replicates.

Next, we assessed how impaired TCR movement and centralization, along with the observed alterations in downregulation and recycling, would affect proximal TCR signalling. To this end, we allowed T cells to interact with SLBs for 15 minutes, followed by fixation, permeabilization, and antibody staining for the phosphorylated forms of Linker for Activation of T cells (pLAT) and Phospholipase C gamma 1 (pPLCγ1). On *mobile* SLBs, pLAT localized to both central and more peripheral regions of the IS, while pPLCγ1 was almost exclusively found in the cSMAC alongside anti-TCR Fab (**Fig. 5A-E**). Notably, on *mixed-mobility* SLBs both proteins underwent a marked shift in localization from central to more peripheral regions of the IS (**Fig. 5A-E**). While overall staining intensities for pLAT and pPLCγ1 across the entire synaptic interface did not significantly differ between conditions, we noted that T cells on *mixed-mobility* SLBs recruited significantly lower levels of anti-TCR Fab (**Fig. 5F-H**). Interestingly, when pLAT and pPLCγ1 staining intensities were normalized to the level of anti-TCR Fab recruitment in individual cells, we observed significantly increased values for pLAT/TCR and pPLCγ1/TCR under *mixed-mobility* conditions (**Fig. 5I & J**), suggesting enhanced signalling activity of TCR within microclusters in the multifocal IS.

**Figure 5:**
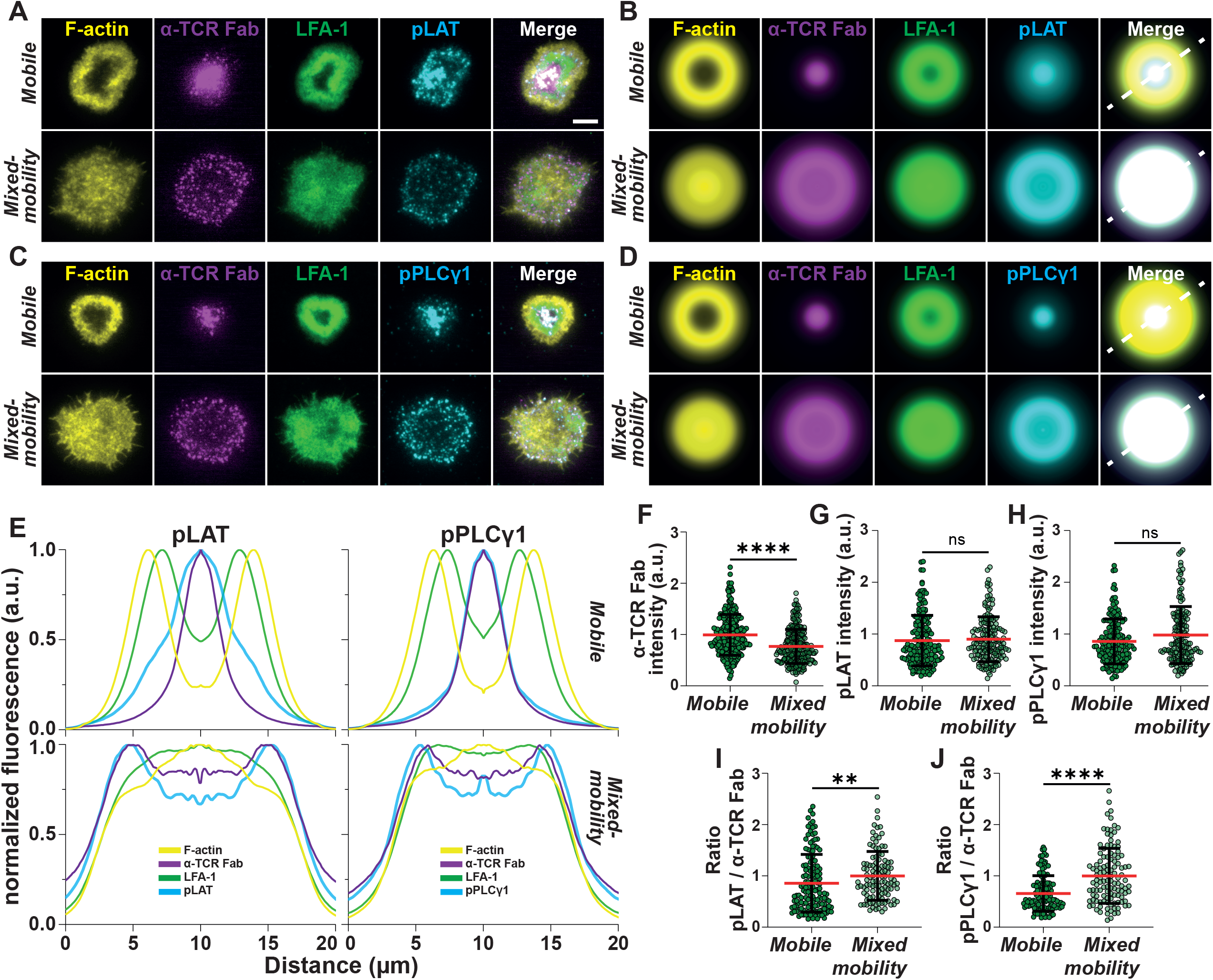
(**A**) Representative TIRF images of synapses formed by T cells on mobile and *mixed-mobility* SLBs, containing fluorescently labelled α-TCR Fab (magenta) after 15min. of incubation. Cells were fixed, stained with a directly antibody for LFA-1 (green), permeabilized and stained with Phalloidin (F-actin, yellow) and an antibody for pLAT or pPLCγ1 (cyan) (**C**) followed by incubation with fluorescently labelled secondary antibodies. Scale bar = 5µm. (**B**) Average intensity projections of radial averages of ≥45 individual synapses formed on mobile or *mixed-mobility* SLBs and stained for pLAT or pPLCγ1 (**D**). (**E**) Normalized intensity profiles along white dashed lines in (B) – (left) and (D) – (right) for all labelled proteins. (**F**) Quantification of normalized α-TCR Fab, pLAT (**G**) and pPLCγ1 (**H**) intensities on mobile and *mixed-mobility* SLBs. Mann-Whitney test, **** < 0.0001. Circles represent measurements from individual cells 3 Biological replicates. (**I**) Quantification of the ratio between α-TCR Fab and pLAT or pPLCγ1 (**J**) respectively. Mann-Whitney test, ** < 0.01, **** < 0.0001. Circles represent measurements from individual cells 3 Biological replicates.

Taken together, these findings suggest that the multifocal IS dampens TCR downregulation through ectocytosis while maintaining EPN1-mediated recycling. In light of our earlier observation that *mixed-mobility* SLBs enhance T cell activation (**Fig. 2A**), this increased response may result from impaired TCR turnover and prolonged surface retention of engaged receptors. Sustained presence of signalling-competent TCRs at the plasma membrane could amplify the activation signal despite changes in the spatial organisation of the IS. This interpretation aligns with previous findings from SLB systems incorporating physical barriers, where TCRs were trapped in the periphery of the synapse and retained phosphotyrosines (pY) for longer than TCRs that translocated to the center of the IS (Mossman et al. 2005). However, then, as here, the effect size was modest.

Next, we assessed the impact of selective ICAM1 immobilization on integrin signalling. Upon engagement of LFA-1, Focal Adhesion Kinase (FAK) is recruited to the site of adhesion, where it undergoes autophosphorylation, creating a docking site for additional signalling molecules. One key downstream effector is Paxillin, a scaffold protein that serves as a platform for the assembly of multi-protein signalling complexes, linking integrin engagement to downstream pathways involved in cytoskeletal remodelling and cellular activation (Harburger and Calderwood 2009). To assess integrin activation, we stained T cells interacting with *mobile* and *mixed-mobility* SLBs for the phosphorylated forms of these proteins (pPaxillin and pFAK). Importantly, we found that *mixed-mobility* SLBs induced significantly higher levels of pFAK (**Fig. 6A & B**) and pPaxillin (**Fig. 6C & D**) compared to *mobile* SLBs. This aligns with earlier findings suggesting that ICAM1, when constrained in its mobility, serves as a more effective ligand for mechanosensitive LFA-1, which requires resistance to the forces exerted by the T cell’s actin cytoskeleton (Comrie et al. 2015; Comrie, Babich, and Burkhardt 2015). This resistance promotes conformational changes in LFA-1 as well as cytoplasmic adapters like CasL (Kumari et al. 2020) that undergoes a conformational change upon cytoskeletal tension-induced stretching, exposing residues that become phosphorylated (pCasL). In support of this hypothesis, T cells on *mixed-mobility* SLBs showed significantly stronger staining for pCasL compared to the *mobile* SLB (**Fig.6E & F**). Notably, these effect sizes were significantly larger than for activating signalling through TCR proximal pathways.

**Figure 6:**
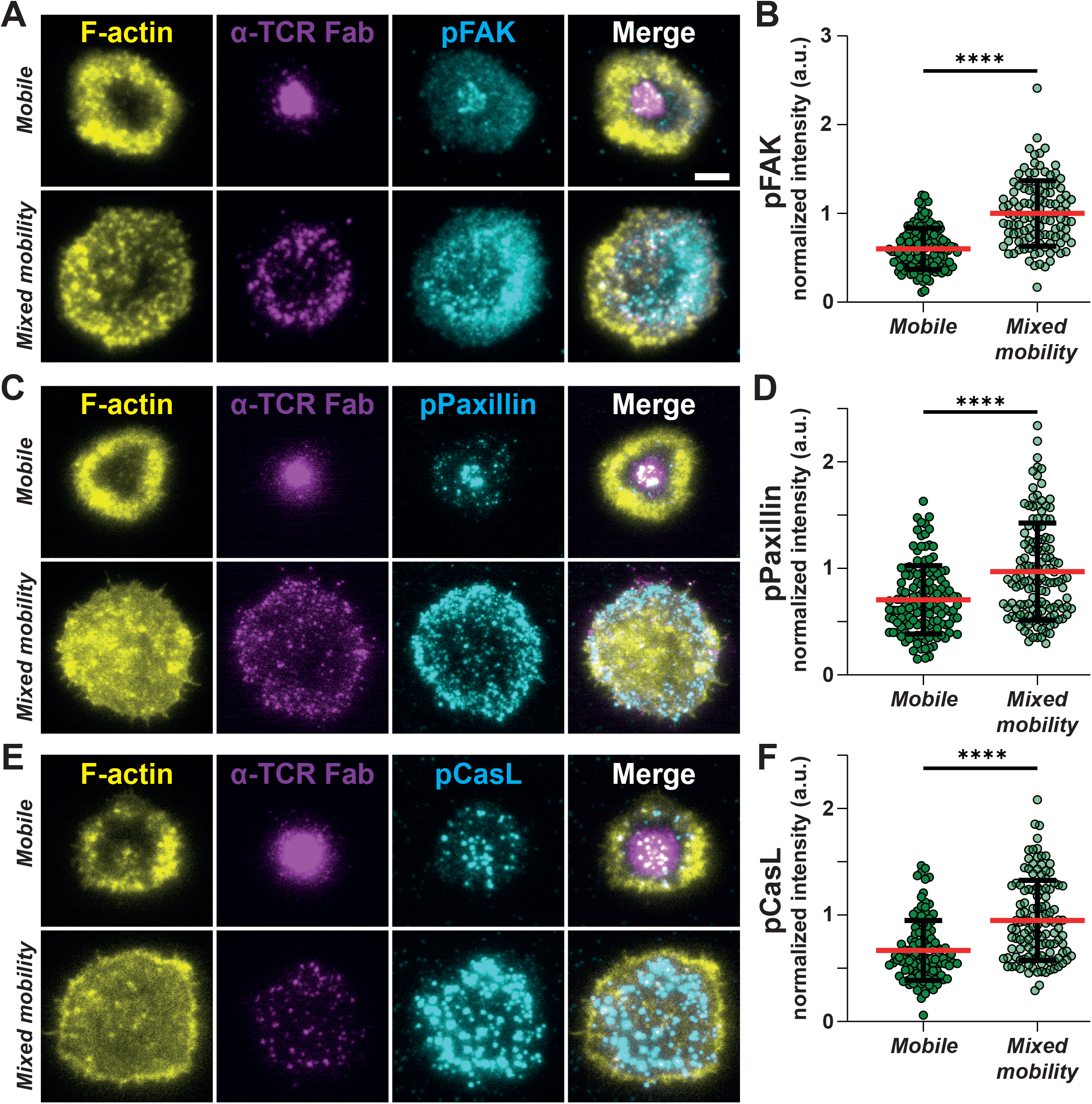
(**A**) Representative TIRF images of synapses formed by T cells on mobile and *mixed-mobility* SLBs, containing fluorescently labelled α-TCR Fab (magenta) after 15min. of incubation. Cells were fixed, permeabilized and stained with Phalloidin (F-actin, yellow) and antibodies for pFAK, pPaxillin (**C**) or pCasL (cyan) (**E**) followed by incubation with fluorescently labelled secondary antibodies. Scale bar = 5µm. (**B**) Quantification of normalized pFAK, pPaxillin (**D**) and pCasL (**F**) intensities on mobile and *mixed-mobility* SLBs. Mann-Whitney test, **** < 0.0001. Circles represent measurements from individual cells. 3 Biological replicates.

We then turned our attention to the effect of ICAM1 immobilization on CD8^+^ T cell activation in a more physiological cell-cell context. To this end, we exploited a newly developed cellular model for immunological studies, termed CombiCells (Patel et al. 2024). CombiCells are CHO cells in which endogenous ICAM1 has been deleted and that express a GPI-anchored version of SpyCatcher (Zakeri et al. 2012), enabling the loading of ligands of choice carrying the SpyTag sequence, in our case CD19. We engineered two new derivatives of CombiCells, one expressing a human ICAM1-FL and another expressing a truncated version of transmembrane ICAM1 lacking the cytoplasmic domain (ICAM1-TL) (Comrie et al. 2015; van Buul et al. 2010). This setup allowed us to assess the impact of ICAM1-FL and ICAM1-TL on T cell-mediated killing across a wide dynamic range of CD19 densities and concentrations of a CD3/CD19 Bispecific T cell engager (BiTE).

Consistent with earlier findings in the SLB system, expression of ICAM1-FL in CombiCells triggered significantly higher levels of IL-2 (**Fig.7A**) and IFN-γ (**Fig.7B**) compared to ICAM1-TL. Importantly, ICAM1-FL also prompted significantly higher levels of cytotoxicity, as measured by endpoint LDH-release assays, compared to ICAM1-TL across all CD19 loading conditions and a wide range of BiTE concentrations (**Fig.7C & D**). Additionally, we performed live-cell imaging of T cell–CombiCell co-cultures in the presence of fluorescent Annexin-V to monitor real-time killing dynamics. In agreement with our endpoint assays, this confirmed that ICAM1-FL induced significantly higher target cell killing compared to ICAM1-TL, while only minimal cell death occurred under control conditions in the absence of CD3/CD19 BiTEs (**Fig.7E & F and Supplementary Movie 3**).

To investigate the mechanistic basis of enhanced cytotoxicity in the context of ICAM1-FL, we seeded T cells on *mobile* and *mixed-mobility* SLBs in the presence of a fluorescently labeled antibody against CD107a, a marker of lytic granules that becomes exposed on the cell surface upon degranulation. After 20 minutes of interaction, followed by fixation, we found that CD8^+^ T cells exhibited significantly higher staining intensity on *mixed-mobility* SLBs compared to *mobile* SLB conditions (**Fig. 7G & H**). In addition, we directly stained for extracellular Perforin-1, a major component of cytotoxic granules that persists as supramolecular attack particles after release into the IS (Bálint et al. 2020). T cells were incubated on SLBs for 5 and 15 minutes before fixation and staining. While both *mobile* and *mixed-mobility* SLBs triggered substantial Perforin-1 release, *mixed-mobility* SLBs induced significantly higher levels at both time points compared to *mobile* SLBs (**Fig. 7I & J**). Notably, lytic granule fusion with the plasma membrane has recently been shown to be associated with mechanically active integrins that experience pulling forces (Wang et al. 2022).

**Figure 7:**
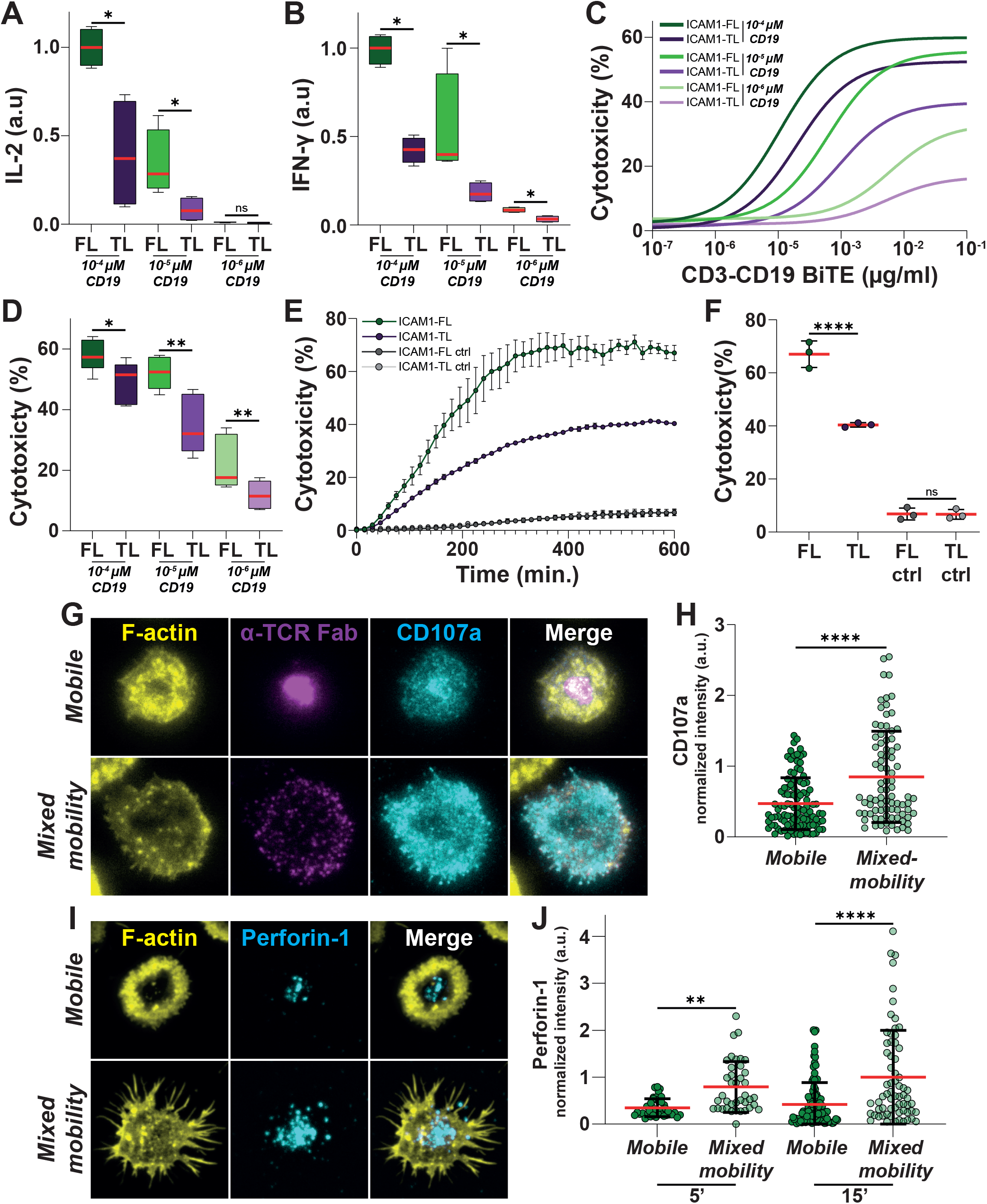
(**A**) Normalized quantification of IL-2 and IFN-γ (**B**) levels in the supernatants of CombiCell-T cell co-cultures in an E:T ratio of 5:1 in the presence of 0.01µg/ml CD3-CD19 BiTE after 24h. Mann-Whitney test, * = 0.0286. 2 Biological replicates. (**C**) Non-linear fits of CombiCell cytotoxicity expressing ICAM1-FL or ICAM1-TL, respectively loaded with different concentrations of CD19 and in the presence of increasing concentrations of CD3-CD19 BiTEs after 18h of co-incubation in an E:T ratio of 5:1. 3 Biological replicates. (**D**) Quantification of cytotoxicity in % at 0.01µg/ml CD3-CD19 BiTE after 18h of co-incubation in an E:T ratio of 5:1. Paired t-test, * = 0.0165, ** ≤ 0.0078. 3 Biological replicates. (**E**) Cytotoxicity over 600min. of ICAM1-FL or ICAM1-TL expressing CombiCells loaded with 0.0001µM of CD19 in co-culture with CD8^+^ T cell in an E:T ratio of 5:1 and in the presence or absence of 0.1µg/ml CD3-CD19 BiTE. Mean ± SEM. 3 Biological replicates. (**F**) Quantification of cytotoxicity after 600min. of co-incubation from (I). One-way ANOVA, **** < 0.0001. (**G**) Representative TIRF images of synapses formed by T cells on mobile and *mixed-mobility* SLBs, containing fluorescently labelled α-TCR Fab (magenta) in the presence of a directly labelled antibody for CD107a (cyan) after 15min. of incubation. Cells were fixed, permeabilized and stained with Phalloidin (F-actin, yellow). (**H**) Quantification of normalized CD107a intensities on mobile and *mixed-mobility* SLBs. Mann-Whitney test, **** < 0.0001. Circles represent measurements from individual cells. 3 Biological replicates. (**I**) Representative TIRF images of synapses formed by T cells on mobile and *mixed-mobility* SLBs, containing fluorescently labelled α-TCR Fab (magenta) after 15min. of incubation. Cells were fixed, stained with a directly labelled antibody for Perforin-1 (cyan), permeabilized and stained with Phalloidin (F-actin, yellow). (**J**) Quantification of normalized Perforin-1 intensities on mobile and *mixed-mobility* SLBs after 5 and 15min. of incubation. Kruskal-Wallis test, ** < 0.0001, **** < 0.0001. Circles represent measurements from individual cells. 2 Biological replicates.

Taken together, these data suggest that immobile ICAM1 is significantly more active than mobile ICAM1 at supporting T cell activation and CD8 T cell effector function.

## Discussion

In this study, we introduce a modular, reductionist SLB-based system that enables the simultaneous presentation of laterally mobile and immobile ligands to T cell-expressed receptors. Building upon previous findings that full-length transmembrane proteins are immobile within SLBs, we demonstrate that reconstitution of full-length ICAM1 into Ni^2+^-NTA-functionalized liposomes with additional His-tagged ligands yields SLBs with spatially interspersed mobile and immobile ligands, enabling controlled manipulation of ligand mobility without altering overall surface composition.

Using this system, we show that selective ICAM1 immobilization enhances CD8^+^ T cell activation in response to mobile anti-TCR Fab fragments, suggesting that physical anchoring of ICAM1 alters downstream cellular behavior and signalling pathways. Mechanistically, our data reveal a disruption of centripetal F-actin flow in the presence of immobilized ICAM1. This aligns with previous reports showing that engagement of surface-immobilized integrin ligands such as ICAM1 or VCAM1, together with co-adsorbed anti-TCR antibodies, attenuates retrograde actin flow (Jankowska et al. 2018; Nguyen, Sylvain, and Bunnell 2008). However, in those systems, the anti-TCR antibodies are themselves surface-immobilized, rendering TCR microclusters immobile regardless of integrin engagement. By contrast, in *mixed-mobility* SLBs, the anti-TCR Fab fragments remain freely mobile and can cluster with the TCR. Strikingly, while cluster formation is preserved, their transport is inhibited: microclusters remain largely at their sites of formation, adopting a multifocal pattern biased towards the periphery of the IS.

These findings connect to the ongoing debate about the role of the cSMAC in TCR signaling. While the cSMAC has historically been viewed as a signaling hub (Grakoui et al. 1999; Monks et al. 1998), and may facilitate signaling at low peptide concentrations by concentrating engaged receptors (Cemerski et al. 2008), multiple lines of evidence suggest it can also serve as a site for signal termination (Varma et al. 2006; Vardhana et al. 2010; Choudhuri et al. 2014). This aligns well with our findings that HRS and the TCR, and later EPN1 and the TCR, are highly co-localized at the center of the IS on *mobile* SLBs, mediating ectocytosis and, respectively, the internalization of the TCR. Furthermore, although overall staining intensities for pLAT and pPLCγ1 did not differ between mobile and *mixed-mobility* SLBs, we found that a considerable fraction of both proteins, particularly pPLCγ1, localized to the cSMAC. Given that PLCγ1 has been detected in synaptic ectosomes of CD4^+^ T cells (Saliba et al. 2019), it is tempting to speculate that a substantial fraction of the central pPLCγ1 signal may be present in extracellular vesicles, and thus, would cease to contribute to cellular signalling.

In contrast, under *mixed-mobility* conditions we find enhanced per-microcluster signaling, as indicated by increased pLAT and pPLCγ1 intensities after normalization for TCR recruitment. This correlates with a shift in TCR trafficking dynamics with a reduction in HRS-mediated ectocytosis while EPN1-dependent TCR internalization is maintained. These findings suggest that the multifocal synapse reduces TCR turnover and prolongs surface retention of signalling competent microclusters, which enhances downstream signalling and increases T cell activation. These findings align with earlier work using SLBs with physical barriers that trap the TCR in the periphery of the IS, generating a small, but significant prolongation of TCR signaling (Mossman et al. 2005). However, it should be noted that in these studies, the transport of both LFA-1/ICAM1 and TCR/pMHC microclusters was blocked, and the effects on integrin signalling were not investigated, an aspect we addressed in this study. We found that, beyond TCR signalling, immobilized ICAM1 enhances integrin signalling. Immobilization promotes stronger recruitment and phosphorylation of FAK and paxillin, in agreement with earlier work showing that mechanical engagement of LFA-1 supports conformational changes in LFA-1 (Comrie, Babich, and Burkhardt 2015; Comrie et al. 2015; Jankowska et al. 2018).

Taken together, our data support models in which the immobilization of integrin ligands promotes T cell activation by attenuating the cytoskeletal transport and the inactivation of microclusters at the center of the IS, thereby enhancing sustained signaling (Nguyen, Sylvain, and Bunnell 2008; Eidell et al. 2021). In addition, integrin signaling itself is enhanced, further contributing to T cell activation. Notably, we also observed increased force transduction. Elevated phosphorylation of CasL, a stretch-sensitive adaptor protein, further supports the notion that immobilized ICAM1 transduces higher cytoskeletal tension, thereby potentiating mechanosensitive downstream signaling pathways.

We did not observe the effects reported in several earlier studies, which likely reflects the specific properties of each experimental system. For example, modulating the mobility of both anti-TCR antibody and ICAM1 by altering the lipid composition of the SLB led to decreased signalling by immobile bilayers (Hsu et al. 2012). Similarly, in a system using glass-bound TCR agonist and VCAM-1, TCR signaling was shown to decrease progressively with increasing amounts of VCAM-1, correlating with a reduction in F-actin flow. A smaller, but significant effect was also observed with high concentrations of ICAM1. These findings have been interpreted to suggest that reduced actin flow diminishes mechanical forces on the TCR, thereby impairing signalling (Jankowska et al. 2018). However, it is important to note that the density of ICAM1 in glass-bound systems is often unclear and may be orders of magnitude higher than in our system. Excessive integrin engagement under such conditions could reduce F-actin dynamics to an extent that mechanical forces on the TCR are effectively lost, leading to reduced signaling. Notably, this interpretation aligns with observations from transgenic mice with constitutively active LFA-1 (Semmrich et al. 2005), or DCs with enhanced coupling of ICAM1 to the cytoskeleton (Leithner et al. 2021), both of which have been associated with dampened T cell activation. In our system, which reproduces the principal state found in natural APCs, where ICAM1 is selectively immobilized while the TCR agonist remains mobile (Comrie et al. 2015), we find that centripetal actin flow is substantially diminished, yet the F-actin cytoskeleton remains highly dynamic and may support both integrin molecular clutch engagement and the generation of mechanical forces on the TCR.

At the level of CD8+ effector functions, ICAM1 immobilization leads to increased lytic granule exocytosis and cytotoxicity by CD8^+^ T cells, both in SLB-based assays and in more physiological co-cultures. Expression of full-length, immobile ICAM1 in target cells enhances T cell-mediated killing and cytokine secretion across a broad range of BiTE and target antigen concentrations. These findings directly support recent reports that mechanically stabilized integrin engagement enhances degranulation (Wang et al., 2022).

More broadly, our system represents an important addition to previous approaches aimed at selectively controlling lateral mobility in SLBs. Existing Polydimethylsiloxane (PDMS)-stamp-based systems generate protein clusters on the micrometer scale and are ultimately limited by the smallest size of stamp that can be generated (Jankowska et al. 2018). In contrast, dSTORM microscopy on *mixed-mobility* SLBs suggests that immobilized proteins are evenly distributed within the SLB, and that protein cluster formation is at the nanometer scale and comparable between mobile controls and immobilized proteins. Another approach, the immobilization of proteins on lithographically nanopatterned surfaces (Cai et al. 2018), allows for nanometer-scale spacing control but requires specialized equipment and expertise in nanopatterning. By comparison, the *mixed-mobility* system presented here can be easily implemented by groups already using SLB technologies to investigate how ligand mobility influences cellular behavior. While we focused on ICAM1, this system could be extended to a wide array of receptors implicated in IS formation, co-stimulation, or checkpoint inhibition.

Finally, our findings raise intriguing questions in the context of tumor immunology. Tumor cells frequently undergo profound cytoskeletal remodeling (Fife, McCarroll, and Kavallaris 2014), which could impact on the lateral mobility of surface proteins, including ligands for T cell-expressed receptors. In this context, drugs targeting the cytoskeleton or its regulatory pathways may inadvertently modulate immune recognition, not by altering ligand expression *per se*, but by modifying mechanical parameters such as mobility and resistance to cytoskeletal tension. A deeper understanding of these processes could contribute to development and refinement of therapeutic interventions. Furthermore, incorporating targeted mobility constraints into synthetic systems for therapeutic applications, such as artificial antigen presenting cells (van der Weijden et al. 2014), may enhance their ability to engage mechanosensitive immune receptors and thereby increase their immunostimulatory potency.

## Acknowledgments

M.L.D. and the Kennedy Institute Strategy was supported by the Kennedy Trust for Rheumatology Research. A.L. was funded by an Erwin Schrödinger postdoctoral fellowship of the Austrian Science Fund (FWF, project number: J4542-B) and was an EMBO non-stipendiary postdoctoral fellow (ALTF 1109-2020). A.L. and M.L.D. are supported by the Biotechnology & Biological Sciences Research Council (BBSRC, BB/X015408/1). T.M. was supported by the CAMS-Oxford Institute Senior Postdoctoral Research Fellowship, funded by the CAMS Innovation Fund for Medical Science (CIFMS), China (Grant: 2018-I2M-2-002). We would like to thank the Oxford-ZEISS Centre of Excellence in Biomedical Imaging for their excellent support.

---

**Supplementary Movie 1:** Time-lapse TIRF microscopy of Lifeact-mCitrine (yellow, top) transfected CD8^+^ T cells making first contact with α-TCR Fab (magenta, bottom) containing mobile (left) or *mixed-mobility* (right) SLBs. Frame interval: 3 sec.

**Supplementary Movie 2:** Time-lapse TIRF microscopy of Lifeact-mCitrine (yellow) transfected CD8^+^ T cells interacting with mobile (left) or *mixed-mobility* (right) SLBs. Frame interval: 3 sec.

**Supplementary Movie 3:** Time-lapse microscopy of CFSE labelled CombiCells (cyan), loaded with 0.0001µM of CD19 and expressing either ICAM1-FL or ICAM1-TL, in co-culture with CD8^+^ T cells in the presence of Annexin-V (magenta), and in the presence or absence of 0.1µg/ml CD3-CD19 BiTE. Frame interval: 15 min.

