## Supplementary material for "Mixed-mobility supported lipid bilayers reveal the modulatory role of immobilized ICAM1 on T cell activation, effector functions and immune synapse organization": Material and Methods

***Cell culture:*** Human primary $CD8^+$ cells and CombiCells were routinely cultured in RPMI (Gibco) medium supplemented with 10% fetal bovine serum, 1% penicillin/streptomycin, 1% L-glutamine, 1% non-essential amino acids and 50 mM HEPES in cell culture treated flasks at 37°C, 5% $CO_2$ and 100% humidity. CombiCells were split at ~80% confluency by trypsinization. For $CD8^+$ culture and expansion, 50 U/mL recombinant human IL-2 (PeproTech, UK) were added to the culture medium. Primary human $CD8^+$ were isolated from healthy human donor blood, provided by the National Health Service (NHS) blood service under ethics agreement number 11/H711/7, with commercially available negative selection kits (RosettaSep Human $CD8^+$ T cell Enrichment Cocktail, STEMCELL technologies, UK) following the manufacturers instruction. After isolation, cells were expanded by addition of anti-CD3/anti-CD28 T cell activation beads (Dynabeads ThermoFisher Scientific, UK) at 25 µL/ $10^6$ cells for 3 days following the manufacturer's instructions. The magnetic beads were subsequently removed and cells were expanded further at $10^6$ cells/mL for an additional 4 days. T cells were either used within 2 days for experiments or cryopreserved.

***Lentivirus production and infection:*** DNA sequences encoding full-length human ICAM1 (ICAM1-FL) and cytoplasmic tail-depleted ICAM1 (ICAM1-TL) were designed based on previously published sequences (van Buul et al. 2010), synthesized and cloned into the pLenti6.3 vector (ThermoFisher). Lentiviruses were produced in HEK293 cells by co-transfecting ICAM1 expression plasmids with the packaging and envelope plasmids pMD2.G (Addgene, #12259) and pΔ8.91 (Addgene, #202687) using the transfection reagent Genejuice (Merck). Supernatants were harvested after 72h, filtered through a 0.45µm filter and either flash frozen or used immediately to infect CombiCells. ICAM1-FL and TL infected CombiCells were expanded for 1 week, after which they were trypsinized, stained for ICAM1 using a fluorescently labelled antibody and sorted for equal surface of expression levels. Sorted ICAM1-FL and TL cells were expanded and frozen in multiple aliquots to perform experiments using cells from low passage numbers. Cultured cells were routinely checked to ensure comparable ICAM1 surface expression levels.

***Formation of mobile and mixed-mobility SLBs:*** The procedure for reconstituting proteins into liposomes was based on previously published protocols (Dustin et al. 2007). In brief, ICAM1 was isolated from human spleen tissue by homogenizing ~8g of tissue in 50ml lysis buffer (25mM Tris.Cl, 0.15M NaCl, 1% Triton X-100, 5mM EDTA, 5 tablets of Roche protease inhibitor cocktail, pH 8 at 4℃) using a glass homogenizer. The lysate was then filtered through a cell strainer, incubated on ice for 1h and ultracentrifuged at 100.000g for 1h at 4℃. The supernatant was collected, filtered through a 0.2µm filter and either frozen at -80℃ or directly used for the affinity isolation of ICAM1.

Affinity-isolation matrices for ICAM1, as well as matrices coupled to irrelevant antibodies, were prepared by resuspending 750mg of NHS-Agarose (26197, Thermofisher) with either 25mg of anti-human ICAM1 antibody (BioXcell, BE0020-2-25MG) or IgGs from mouse serum (I5381, Merck) in 10ml of Coupling/Wash buffer (0.1M sodium phosphate, 0.15M NaCl, pH 7.2) followed by end-over-end mixer at 4℃ overnight. Agarose matrices were centrifuged for 1min. at 1000g and washed twice with Coupling/Wash buffer, followed by incubation with Quenching buffer (1M Ethanolamine, pH 7.4) for 15min. and two additional washes with Coupling/Wash buffer. Matrices were either stored in PBS at 4℃ or directly used for the isolation of ICAM1 from tissue lysates.

For affinity isolation, 2.5ml of matrices coupled to irrelevant mouse IgGs or 1.5ml of anti-ICAM1 functionalized matrices were transferred to 15ml tubes using wide-bore tips. To remove loosely bound antibodies, matrices were washed 3x with low-pH buffer (50mM Glycine, 0.15M NaCl, 1% Triton X-100, pH 3), 3x with high-pH buffer (50mM Triethanolamine, 0.15M NaCl, 1% Triton X-100, pH 12) and 3x with Wash buffer 1 (25mM Tris.Cl, 0.15M NaCl, 1% Triton X-100, pH 8 at 4℃). Lysates were cleared by two sequential incubations with 1ml of irrelevant IgG-coupled agarose matrices for 1h each at 4℃ with end-over-end mixing. Cleared lysates were then incubated overnight at 4℃ with 1.5ml of anti-ICAM1 functionalized agarose matrices, again with end-over-end mixing. Agarose matrices were washed 3x with Wash buffer 1 and 3x with Wash buffer 2 (25mM Tris.Cl, 0.15M NaCl, 1% Octyl-β-glucoside, pH 8 at 4℃) and transferred to a Poly-prep chromatography column (731-1550, Biorad). ICAM1 was eluted by adding 2ml of low-pH elution buffer (50mM Glycine, 0.15M NaCl, 1% Octyl-β-glucoside, pH 3) to the column, collecting the eluate into 200µl of neutralisation buffer (1M Tris.Cl, 1% Octyl-β-glucoside, pH 8.5). Aliquots of eluate were loaded onto SDS-PAGE for Coomassie staining and Western plotting to confirm the isolation of ICAM1.

For reconstitution of ICAM1 into liposomes, a 20x stock solution containing 2.5% DOGS-NTA(Ni2+) and 97.5% DOPC (all lipid reagents were obtained from Avanti Polar Lipids) at a concentration of 4mM was prepared by mixing appropriate amounts of lipid in chloroform, followed by vacuum desiccation and resuspension into Tris-saline buffer (25mM Tris.Cl, 0.15M NaCl, pH 8 at 4℃) containing 2% Octyl-β-glucoside. The ICAM1-containing solution was then mixed with this stock solution to achieve a final 1x concentration. Control liposomes without ICAM1 were prepared in the same way, using buffer instead of ICAM1-containing solution. These solutions were transferred to dialysis tubes (GeBaflex maxi, 8kDa MWCO) and dialyzed for 48h against Tris-saline, with 4x buffer exchanges.

For the formation of Glass-supported lipid bilayers, plasma cleaned glass cover slips (SCHOTT UK) were fixed to 6-channel flow chambers (sticky-Slide VI 0.4, Ibidi). Channels were filled with either ICAM1-containing proteoliposome solutions or control liposomes, incubated for 20min., washed twice with flow buffer (20mM HEPES, 1.37mM NaCl, 5mM KCl, 0.5mM $Na_2HPO_4$, 6mM D-Glucose, 1mM $CaCl_2$, 2mM $MgCl_2$, 1% BSA, pH 7.2) and blocked with 3% BSA in flow buffer for 20min.. This was followed by two additional washes with flow buffer and subsequent incubation with either, His-tagged and Alexa-647 labelled α-TCR Fab in the case of ICAM1-containing proteolipsomes to obtain *mixed-mobility* SLBs, or in the case of control liposomes with His-tagged α-TCR Fab and the His-tagged extracellular domain of ICAM1 to obtain mobile SLBs. Specific protein concentrations required to achieve a molecular density of 30 molecules/$\mu m^2$ for α-TCR Fab and, for His-tagged ICAM1, to match the ICAM1 density in proteoliposomes, which ranged from ~150 - 300 molecules/$\mu m^2$, were determined from flow-cytometry-based calibration experiments on bead-supported lipid bilayers compared to fluorescently-labelled reference beads of known molecular densities (Bangs Laboratories).

***FRAP and FCS:*** For photobleaching experiments, mobile and *mixed-mobility* SLBs were stained with an Alexa-488 labelled anti-ICAM1 antibody (HCD54, BioLegend), washed 3x with flow buffer and transferred to a Zeiss LSM980 confocal microscope, equipped with a C-Plan Apo 63x 1.40NA oil immersion objective and using Zeiss 518F23C immersion oil. Before photobleaching, SLBs were imaged for 1sec. at randomly selected regions, bleached and imaged for another 60sec. Adjacent, non-bleached regions served as controls for bleaching corrections. Spot FCS experiments were performed on a Zeiss LSM980 with a Plan Apo 40x 1.20NA water immersion objective and using distilled H2O as immersion liquid. These experiments were performed with a pinhole diameter of 37um (corresponding to 1 Airy unit at 488 nm) and using internal GaAsP point detectors. These experiments were performed with a laser power setting of 1% of the 488 nm laser and 2% of the 640 nm laser such that we observed no photobleaching during the acquisition.

Data analyses were performed using Fiji ImageJ, FoCuS-point (Waithe et al. 2016) and Graph Pad prism.

***dSTORM imaging:*** The Oxford NanoImager (ONI) microscope was used to generate 2D SMLM images using dSTORM. A 100x objective lens/1.4 NA in oil-immersion was used. 405 nm and 488 nm lasers were used with 2 channels (with dichroic mirror split at 640 nm). Mobile and *mixed-mobility* SLBs were stained with an Alexa-488 labelled anti-ICAM1 antibody as described above. All calibrations and images were performed at 32°C with a supercritical TIRF angle of 51.5° and paired samples acquired with the exact same preparation of dSTORM buffer (320 µl 1x PBS, 40 µl 50% glucose, 40 µl cysteamine 688 (MEA), 4 µl glucose-oxidase).

The 488 nm laser was used at 60% laser power to excite anti-ICAM1-Alexa-488. The 405 nm laser was used to promote fluorophore blinking. 8000 frames with an exposure of 30ms were acquired.

The Oxford NanoImager (ONI) analysis platform, CODI, was used to analyse dSTORM images. Constrained clustering algorithm included in CODI was used to identify clusters. Drift correction was applied using the DME algorithm. Frames were filtered first according to acquisition program settings then to exclude the initial excitation peaks for each laser. The steady state of localizations per frame was included as true blinking localizations. Filtering parameters around the sigma peak were then applied to include a sigma range of ~50-250 nm (Gaussian fit to localize single molecules). Localization precision filtering was then applied to include localization with precision less than or equal to 25 nm. ICAM-1 events were depicted with a display sigma of 5 nm using fixed representation in the representative images, which were downloaded from CODI in TIFF and PNG formats then cropped and converted to PNG micrographs for figures.

***Immunofluorescence and live imaging:*** T cells were added to supported lipid bilayers (SLBs) in flow buffer at 37 °C with 5% $CO_2$ for the indicated time periods. Fixation was then performed by adding paraformaldehyde to a final concentration of 4% and incubating for 15min. at 37 °C. For extracellular staining, cells were blocked with 5% BSA in PBS for 1h and incubated with directly labeled primary antibodies (anti-CD11a-AlexaFluor 405 (TS2/4), anti-CD107a-AlexaFluor 647 (H4A3), and anti-Perforin-AlexaFluor 647 (B-D48)) for 1h at a 1:100 dilution. For intracellular stainings, cells were permeabilized with 0.1% Triton X-100 for min., followed by PBS washes and blocking with 5% BSA in PBS for 1h. Directly labeled primary antibodies (anti-β-Tubulin-AlexaFluor 488 (9F3), 1:200; anti-HRS-AlexaFluor 568, Abcam #72053, 1:200; anti-Epsin 1- AlexaFluor 647 (EPR3023), 1:200) or unlabeled primary antibodies (anti-pLAT, Cell Signaling #3581; anti-pPLCγ1, Cell Signaling #2821; anti-pFAK (31H5L17); anti-pPaxillin (E9U9F); anti-pCasL, Cell Signaling #4015) were added at a 1:100 dilution unless otherwise indicated and incubated at 4°C overnight. This was followed by incubation with appropriate fluorescently labeled secondary antibodies and phalloidin to visualize F-actin. For confocal microscopy cells were imaged with a Zeiss LSM980 equipped with an AiryScan module and a Plan-Achromat 63X oil immersion objective NA 1.4. TIRF microscopy was performed on an Olympus IX83 microscope equipped with 405, 488, 568 and 640nm laser lines, a 100X oil immersion objective NA 1.45 and an EMCCD camera.

***Cell-cell assays:*** All functional assays involving primary human $CD8^+$ T-cell/CombiCell co-cultures were performed in 96-well transparent flat bottom well plate formats (Costar 96) with a total culture volume of 150 µL and a CombiCell to T cell ratio of 1:5 with a co-incubation time of 18 hours. CombiCells were seeded one day prior to the experiment in 100 µL fully-supplemented culture medium at a concentration of 10.000 cells/well to allow adhesion. On the day of the experiment, CombiCells were loaded with the indicated CD19-SpyTag concentrations for 1h, washed three times with fresh cell culture media, before 50.000 $CD8^+$ T-cells were added for a total volume of 125 µL per well. Dilution series of anti-CD3/CD19 BiTE (Invivogen) were prepared in 25µl per well, resulting in a final co-culture volume of 150 µl per well and the indicated final BiTE concentrations. LDH release assays (CyQuant LDH cytotoxicity assay, Thermo Fischer, USA) and cytokine measurements (BD Cytometric Bead Array (CBA) Human Th1/Th2 Cytokine Cytometric Bead Array Kit) were performed from cell culture supernatants, following the manufacturers instructions.

Live monitoring of cytotoxicity was achieved by staining CombiCells with 20µM CFSE, followed by seeding into the wells of a 18-well slide (iBidi) at 20.000 cells/well and incubation overnight at 37°C, 5% $CO_2$. CombiCells were loaded with 0.0001µM CD19-SpyTag, washed and T cells were added at 100.000 cells/well in the presence of 0.1µg/ml anti-CD3/CD19 BiTE and 1/200 Red Incucyte Annexin-V dye (Sartorius). Imaging was performed on a LSM980 confocal microscope equipped with an absolute focus module in multi-position mode. Images were collected every 30 min. for 12h. Image analysis was performed by segmenting the CFSE and Annexin-V signals using machine learning based image segmentation software ilastik, followed by calculation of the relative Annexin-V positive area in ImageJ.

Buul, Jaap D. van, Jos van Rijssel, Floris P. J. van Alphen, Mark Hoogenboezem, Simon Tol, Kees A. Hoeben, Jan van Marle, Erik P. J. Mul, and Peter L. Hordijk. 2010. “Inside-out Regulation of ICAM-1 Dynamics in TNF-Alpha-Activated Endothelium.” *PloS One* 5 (6): e11336.

Dustin, Michael L., Toby Starr, Rajat Varma, and V. Kaye Thomas. 2007. “Supported Planar Bilayers for Study of the Immunological Synapse.” *Current Protocols in Immunology* Chapter 18 (February):18.13.1–18.13.35.

Waithe, Dominic, Mathias P. Clausen, Erdinc Sezgin, and Christian Eggeling. 2016. “FoCuS-Point: Software for STED Fluorescence Correlation and Time-Gated Single Photon Counting.” *Bioinformatics (Oxford, England)* 32 (6): 958–60.
